# E3 ubiquitin ligase HUWE1 mediates HBx-dependent K6-linked polyubiquitylation and stabilization of Nrf2 to restrict hepatitis B virus replication

**DOI:** 10.64898/2026.04.20.719611

**Authors:** Muchamad Ridotu Solichin, Lin Deng, Hifdza Faza Felisha, Yeshua Putra Krisnugraha, Chieko Matsui, Takayuki Abe, Akihide Ryo, Koichi Watashi, Masamichi Muramatsu, Ikuo Shoji

## Abstract

Hepatitis B virus (HBV) infection remains a major global health burden, and HBV X protein (HBx) plays a central role in modulating host pathways that influence viral replication. We previously reported that the oxidative stress sensor Kelch-like ECH-associated protein 1 (Keap1) recognizes HBx to activate the NF-E2-related factor 2 (Nrf2) signaling pathway to suppress HBV replication. Although canonical K48-linked ubiquitylation is known to control Nrf2 turnover, the contribution of non-canonical ubiquitin linkages to Nrf2 regulation during HBV infection remains unclear. Here, we investigated the role of HECT, UBA, and WWE domain-containing E3 ubiquitin ligase 1 (HUWE1) in the regulation of Nrf2 in the context of HBV replication. Cell-based ubiquitylation assays demonstrated that HUWE1 knockdown reduced HBx-mediated K6-linked polyubiquitylation of Nrf2, while overexpression of wild-type HUWE1, but not the catalytically inactive HUWE1(C4341A) mutant, enhanced it. Coimmunoprecipitation and proximity ligation assays demonstrated that HUWE1 interacts with HBx in the cytoplasm and binds Nrf2 only in the presence of HBx, suggesting that HBx promotes the interaction between HUWE1 and Nrf2. Cycloheximide chase assays demonstrated that HUWE1 knockdown destabilized Nrf2 in HBx-expressing cells. Furthermore, depletion or pharmacological inhibition of HUWE1 increased intracellular HBV RNA and pgRNA levels as well as extracellular HBV DNA and HBsAg levels in HBV-infected cells. Collectively, these results support a model in which HUWE1 mediates HBx-dependent K6-linked polyubiquitylation and stabilization of Nrf2 to restrict HBV replication. This study expands current understanding of non-canonical ubiquitin signaling in HBV-host interactions.

**DATA SUMMARY:** All data are presented in the main figures. The data that support the findings of this study is available at bioRxiv (https://doi.org/10.64898/2026.04.20.719611). Raw sequencing data, microscopy images, materials, and sequence information are available upon request. Correspondence and requests for materials should be addressed to Professor Ikuo Shoji.

**IMPACT STATEMENT:** Hepatitis B virus (HBV) chronically infects approximately 254 million people worldwide, yet host mechanisms that restrict viral replication remain incompletely understood. The Keap1/ Nrf2 signaling pathway is a central defense against oxidative stress. Under basal conditions, Nrf2 is targeted for degradation via Keap1/Cullin3-mediated K48-linked polyubiquitylation. Here, we provide evidence that the E3 ubiquitin ligase HUWE1 contributes to HBx-dependent K6-linked polyubiquitylation and stabilization of Nrf2. Our findings support a model in which non-canonical ubiquitin signaling helps shape the HBV-host interactions and contributes to suppression of viral replication. This study extends current understanding of the ubiquitin code in HBV infection and highlights HUWE1 as a candidate component of an anti-HBV regulatory pathway.

## INTRODUCTION

Chronic infection with hepatitis B virus (HBV) remains a major global health challenge. The World Health Organization estimates that 254 million people have chronic HBV infection worldwide, and complications such as HBV-related cirrhosis and hepatocellular carcinoma were responsible for approximately 1.1 million deaths in 2022 [1, 2]. Current treatments include nucleos(t)ide analogues and interferon-α, but they rarely achieve a functional cure, underscoring the urgent need for novel therapeutic strategies [3].

HBV is a hepatotropic virus composed of a 3.2-kb partially double-stranded relaxed circular DNA (rcDNA) genome [4]. In infected hepatocytes, rcDNA is converted to covalently closed circular DNA (cccDNA), which serves as a template for the production of viral proteins, including HBV surface antigen (HBsAg), HBV core antigen (HBcAg), HBV e antigen (HBeAg), HBV polymerase (Pol), and HBV X protein (HBx) [4, 5]. Among these viral proteins, HBx is a multifunctional regulatory protein that is essential for initiating cccDNA transcription and sustaining HBV replication, and has also been associated with hepatocarcinogenesis [5, 6]. The Kelch-like ECH-associated protein 1 (Keap1)/NF-E2-related factor 2 (Nrf2) signaling pathway is a major defense mechanism against oxidative stress [7]. Under basal conditions, Keap1 binds Nrf2 in the cytoplasm and promotes Nrf2 polyubiquitylation and degradation via K48-linked ubiquitin chains [8]. However, under oxidative stress conditions, Keap1 becomes inactivated, leading to Nrf2 stabilization. Nrf2 subsequently translocates to the nucleus and activates transcription of antioxidant response genes [8–10].

We previously reported that during HBV infection, Keap1 recognizes the HBx protein, resulting in Nrf2 stabilization and activation. Activated Nrf2 then translocates to the nucleus and inhibits HBV replication by suppressing HBV promoter activity [11]. Interestingly, we also found evidence that HBx promotes non-canonical polyubiquitylation of Nrf2, including K6-linked chains, and that knockdown of HECT, UBA, and WWE domain-containing E3 ubiquitin ligase 1 (HUWE1) reduced HBx-mediated K6-linked polyubiquitylation of Nrf2 [11]. While canonical K48-linked ubiquitylation mediated by the Keap1-Cullin3 complex is known to regulate Nrf2 degradation [8, 10], it remains unclear whether non-canonical ubiquitin linkages, particularly K6-linked chains, contribute to Nrf2 regulation during HBV infection.

HUWE1 is a 482-kDa E3 ubiquitin ligase belonging to the HECT family of ubiquitin ligases, also known as Mule [12], ARF-BP1 [13], HECTH9 [14], URE-B1 [15], LASU1, or E3Histone [16]. It was first identified as an enzyme that promotes the degradation of several key regulators of stress response and survival pathways, including Mcl-1, p53, and c-Myc [12–14]. In addition to the C-terminal HECT domain, HUWE1 contains multiple functional domains, including a ubiquitin-associated domain, a Bcl-2 homology region 3 domain, and a UBM1 domain. HUWE1 catalyzes K48-, K63-, and K6-linked ubiquitin conjugation [17]. Importantly, HUWE1 has been reported to be a major E3 ligase responsible for generating non-canonical K6-linked ubiquitin chains [18]. Unlike the more widely recognized K48- or K63-linked chains, which are associated with protein degradation or signaling processes, K6-linked polyubiquitylation has recently attracted attention for its role in non-degradative functions such as protein stabilization [19, 20]. HUWE1 possesses a catalytic cysteine at C4341 in the HECT domain [21].

Here, we investigated whether HUWE1 contributes to HBx-dependent regulation of Nrf2 and examined the consequences of this pathway for HBV replication. Our findings suggest that HUWE1 promotes HBx-dependent K6-linked polyubiquitylation and stabilization of Nrf2 and thereby contributes to the restriction of HBV replication.

## METHODS

### Cell culture

Human hepatoblastoma HepG2 cells were maintained in Eagle’s minimum essential medium (EMEM) containing L-glutamine (FUJIFILM Wako Pure Chemical Industries, Osaka, Japan) and supplemented with 10% heat-inactivated fetal bovine serum (FBS) (Biowest, Nuaillé, France), 100 units/mL penicillin, 100 μg/mL streptomycin (Gibco, Grand Island, NY, USA), and 0.1 mM nonessential amino acids (Gibco). HepG2-NTCP-Myc-His_6_ cells stably expressing sodium taurocholate cotransporting polypeptide (NTCP) [22] were maintained in Dulbecco’s modified Eagle’s medium (DMEM; high glucose) with L-glutamine (FUJIFILM Wako Pure Chemical Industries) supplemented with 10% FBS (Biowest), 10 mM HEPES (Gibco), 100 units/mL penicillin, 100 μg/mL streptomycin (Gibco), 0.1 mM nonessential amino acids (Gibco), and 1 ng/mL puromycin (Sigma-Aldrich, St. Louis, MO, USA). PXB primary human hepatocytes (PhoenixBio, Hiroshima, Japan) were maintained in dHCGM medium supplied by the manufacturer. All transfections were performed using FuGene 6 transfection reagent (Promega, Madison, WI, USA) following the manufacturer’s protocol.

### HBV preparation and infection

HBV (genotype D, subtype ayw) was prepared from the culture supernatant of Hep38.7-Tet cells as described previously [22]. Briefly, the supernatants were clarified by centrifugation, then HBV particles were precipitated overnight at 4°C with 12.5% (wt/vol) polyethylene glycol 6000 (PEG-6000) (Hampton Research, Aliso Viejo, CA, USA) and 0.75 M NaCl. The pellet was collected by centrifugation at 9,100 × g for 60 min at 4°C, resuspended, and further purified through a 20% sucrose cushion by ultracentrifugation at 110,000 × g for 16 h at 4°C using an SW41 Ti rotor (Beckman Coulter, Brea, CA, USA). The purified HBV was resuspended in Opti-MEM (Thermo Fisher Scientific, Waltham, MA, USA) and stored at -80°C. HBV genome equivalents (GEq) were quantified by quantitative PCR (qPCR). HBV plasmid DNA pUC19-HBV-D-IND60 was used as the standard. HBV infection was performed as described previously [23]. HepG2-NTCP-Myc-His_6_ cells or PXB cells were inoculated with HBV at 1,000 GEq/cell in the presence of 4% PEG-8000 at 37 °C for 24 h, then washed twice with medium and cultured in fresh medium.

### HBV NanoLuc assay

The HBV NanoLuc (HBV/NL) encodes NanoLuc luciferase in place of part of the preCore open reading frame [22, 24]. HepG2-NTCP-Myc-His_6_ cells were inoculated with HBV/NL at 100 GEq/cell. At 4 days postinfection (dpi), cells were lysed and NanoLuc luciferase activity was measured using the Nano-Glo Luciferase Assay Reagent (Promega) and a GloMax Navigator microplate luminometer GM2000 (Promega).

### Expression plasmids

The expression plasmid pDEST-FLAG-HUWE1 was purchased from Addgene (Watertown, MA, USA). To construct pCAG-HUWE1 and pCAG-HA-HUWE1, the cDNA fragment of HUWE1 was amplified by PCR using pDEST-FLAG-HUWE1 as a template. The primer sequences were as follows: forward primer (untagged HUWE1), 5’-AGAGGTACCGCGGCCATGAAAGTAGACAGGACTAAACTG-3’; reverse primer (untagged HUWE1), 5’-GGCCTGCAGGCGGCCTTAGGCCAGCCCAAAGCC-3’; forward primer (HA-HUWE1), 5’-TCGAGCTCAGCGGCCATGAAAGTAGACAGGACTAAACTG-3’; and reverse primer (HA-HUWE1), 5’-AGTGAATTCGCGGCCTTAGGCCAGCCCAAAGCC-3’. The amplified PCR products were purified and cloned into the NotI site of pCAG-MSC2 or pCAG-HA, respectively, using In-Fusion HD cloning kit (Clontech, Mountain View, CA, USA). The catalytically inactive mutant pCAG-HUWE1(C4341A) was generated by site-directed mutagenesis using the following primers: forward primer, 5’-GCCTGCCTTCAGCTCACACAGCTTTTAATCAGCTGGATCT-3’; reverse primer, 5’-AGATCCAGCTGATTAAAAGCTGTGTGAGCTGAAGGCAGGC-3’.

The siRNA-resistant rescue plasmid pCAG-HUWE1_R was generated by site-directed mutagenesis using pCAG-HUWE1 as a template. The primer sequences for PCR were as follows: forward primer, 5’-CTCCGAAGCCCGGCTTTCACAAGCCGCTTAAGTGGCAACC-3’; reverse primer, 5’-GGTTGCCACTTAAGCGGCTTGTGAAAGCCGGGCTTCGGAG-3’. The sequences of the constructed plasmids in this study were confirmed by DNA sequencing (Eurofins Genomics, Tokyo, Japan).

The HBV genome plasmids pUC19-HBV-C-AT_JPN and pUC19-D-IND60, the HBx-deficient derivative plasmid pUC19-HBV-C-AT_JPN(ΔHBx), pEF1A-HBx-Myc-His_6_, pEF1A-HBc-Myc-His_6_, pEGFP-C3-HBx, pCAG-FLAG-Nrf2, pRK5-HA-Ub (wild-type), and pRK5-HA-Ub-K6 were described previously [11, 25–27].

### Antibodies and reagents

The mouse monoclonal antibodies (MAbs) used in this study were anti-Nrf2 MAb (A-10; sc-365949; Santa Cruz Biotechnology, Santa Cruz, CA, USA), anti-FLAG (M2) (F3165; Sigma-Aldrich), anti-c-Myc MAb (9E10; sc-40; Santa Cruz Biotechnology), anti-HBc MAb (clone 7B2, hybridoma culture supernatant) [28], and anti-glyceraldehyde-3-phosphate dehydrogenase (GAPDH) MAb (014-25524; FUJIFILM Wako Pure Chemical Industries). The rabbit polyclonal antibodies (PAbs) used in this study were anti-HUWE1 PAb (ab70161; Abcam, Cambridge, UK), anti-HA PAb (H-6908; Sigma-Aldrich), anti-FLAG PAb (2368; Cell Signaling Technology, Danvers, MA, USA), and anti-HBx PAb (ab39716; Abcam). Horseradish peroxidase (HRP)-conjugated anti-mouse IgG (7076; Cell Signaling Technology) and HRP-conjugated anti-rabbit IgG (7074; Cell Signaling Technology) were used as secondary antibodies. The HUWE1 inhibitor BI8626 (TargetMol Chemicals, Wellesley Hills, MA, USA) was dissolved in dimethyl sulfoxide (DMSO).

### Immunoprecipitation

Cells were lysed with the lysis buffer containing 120 mM NaCl, 50 mM HEPES (pH 7.2), 1 mM EDTA (pH 8.0), 1% NP-40, 0.5% sodium deoxycholic acid, and protease inhibitor cocktail (Roche, Mannheim, Germany) on ice for 30 min. The lysates were cleared by centrifugation at 20,400 × g for 15 min at 4°C. The supernatant was subjected to immunoprecipitation with anti-FLAG M2 affinity gel (Sigma-Aldrich) or protein A-Sepharose 4 Fast Flow (GE17-5280-04; GE Healthcare, Buckinghamshire, UK) pre-coupled to the indicated antibody overnight at 4°C with rotation. Beads were washed five times with lysis buffer and the bound proteins were eluted by boiling in SDS sample buffer.

### Immunoblot analysis

Immunoblot analysis was performed as described previously [29]. Proteins were separated by SDS-polyacrylamide gel electrophoresis (SDS-PAGE) and transferred to a polyvinylidene difluoride membrane (Millipore, Billerica, MA, USA). The membranes were incubated with primary antibodies, followed by HRP-conjugated secondary antibodies. The protein bands were visualized using enhanced chemiluminescence western blotting detection reagents (ECL; GE Healthcare). The band intensities were quantified using ImageJ software version 1.56d.

### Cell-based ubiquitylation assay

Cell-based ubiquitylation assays were performed as described previously [30, 31]. HepG2 cells were co-transfected with the indicated plasmids. At 48 h posttransfection, cells were lysed with lysis buffer containing 120 mM NaCl, 50 mM HEPES (pH 7.2), 1 mM EDTA (pH 8.0), 1% NP-40, 0.5% sodium deoxycholic acid, 10 mM *N-*ethylmaleimide (Sigma-Aldrich), and the protease inhibitor cocktail (Roche) for 30 min on ice. The lysates were centrifuged at 20,400 × g for 10 min at 4°C. Cell lysates were denatured by adding SDS to a final concentration of 1% and boiled at 100°C for 10 min and diluted 1:10 in lysis buffer. The diluted lysates were subjected to immunoprecipitation with anti-FLAG M2 affinity beads overnight at 4°C. After being washed, bound proteins were eluted and analyzed by immunoblotting with anti-HA PAb to detect ubiquitylated FLAG-Nrf2.

### siRNA transfection

HepG2, HepG2-NTCP-Myc-His_6_, or PXB cells were transfected with HUWE1 siRNA (SI00757855; Qiagen, Valencia, CA, USA) at a final concentration of 40 nM using Lipofectamine RNAiMAX (Thermo Fisher Scientific) according to the manufacturer’s instructions. AllStars Negative Control siRNA (Qiagen) was used as a control. Except for PXB experiments, all siRNAs were transfected once.

### CHX-chase experiment

To examine the half-life of Nrf2, HepG2 cells were transfected with pCAG-FLAG-Nrf2. At 48 h posttransfection, cells were treated with 100 μg/mL cycloheximide (CHX) (Sigma-Aldrich).Cells designated as the zero-time point were harvested immediately after CHX treatment. For subsequent time points, cells were incubated in CHX-containing medium at 37°C for 1, 2, 4, and 6 h. CHX-treated cells were then subjected to immunoblot analysis. FLAG-Nrf2 band intensities were normalized to GAPDH and expressed as a percentage of the 0-h time point.

### RNA extraction and quantitative reverse transcription-PCR (RT-qPCR)

Total RNA was extracted using the ReliaPrep RNA Cell Miniprep System (Promega) following the manufacturer’s instructions, and cDNA was synthesized using GoScript Reverse Transcription (Promega) with random hexamer primers. qPCR was performed using TB Green Premix Ex Taq II (TaKaRa Bio, Kyoto, Japan) with SYBR green chemistry on a StepOnePlus real-time PCR System (Applied Biosystems, Foster City, CA, USA). The primer sequences were as follows: HBV total RNA (amplifying all HBV transcripts except the 0.8-kb transcript encoding HBx) [32], forward, 5’-GCTTTCACTTTCTCGCCAAC-3’; reverse, 5’-GAGTTCCGCAGTATGGATCG-3’; HBV pgRNA [33], forward, 5’-ACTGTTCAAGCCTCCAAGCTGT-3’; reverse, 5’-GAAGGCAAAAACGAGAGTAACTCCAC-3’; HUWE1 [34] , forward, 5′-CGGCTTTTCCTGAAGAAGGGAC-3′; reverse, 5′-GTTTCCAAAGGCTTCAGAGCAGC-3′.

Human GAPDH gene expression levels were measured as an internal control, using the primers 5’-GCCATCAATGACCCCTTCATT-3’ and 5’-TCTCGCTCCTGGAAGATGG -3’.

### Quantification of extracellular HBV DNA

Extracellular HBV DNA was extracted from culture supernatants using a QIAamp DNA mini kit (Qiagen). HBV DNA was quantified by qPCR using the following primers: forward, 5′-ACCAATCGCCAGTCAGGAAG-3′; and reverse, 5′-ACCAGCAGGGAAATACAGGC-3′. Serial dilutions of plasmid pUC19-HBV-D-IND60 were used as standards.

### Extracellular HBsAg quantification (ELISA)

Culture supernatants were clarified by centrifugation at 1,000 x g for 5 min at 4°C. Extracellular HBsAg levels were quantified using an HBs ELISA kit (DIAsource, Louvain-la-Neuve, Belgium) according to the manufacturer’s protocol.

### Indirect immunofluorescence

Indirect immunofluorescence assay was performed as described previously [29]. In brief, HepG2 cells grown on glass coverslips placed in a 24-well plate were fixed with 4% paraformaldehyde for 15 min at room temperature, permeabilized with 0.1% Triton X-100 in PBS for 15 min at room temperature, and blocked with 1% bovine serum albumin (BSA) in PBS for 1 h. After being washed twice with PBS, the cells were incubated with the primary antibodies followed by the secondary antibodies. The primary antibodies used were anti-c-Myc mouse MAb and anti-FLAG rabbit PAb. The secondary antibodies used were Alexa Fluor 594-conjugated goat anti-mouse IgG (A11005; Molecular Probes, Eugene, OR, USA) and Alexa Fluor 488-conjugated goat anti-rabbit IgG (A11008; Molecular Probes). The stained cells were counterstained with Hoechst 33342 (Molecular Probes) and observed using a confocal laser scanning microscope (LSM700; Carl Zeiss, Oberkochen, Germany).

### Proximity ligation assay (PLA)

In situ PLA was performed using the Duolink In Situ PLA kit (Sigma-Aldrich) as described previously [25]. To evaluate the interaction between HUWE1 and HBx, HepG2 cells cultured on glass coverslips placed in a 24-well plate were co-transfected with pEF1A-HBx-Myc-His_6_ and pCAG-HA-HUWE1. At 48 h posttransfection, the cells were fixed with 4% paraformaldehyde for 15 min and permeabilized with PBS containing 0.1% Triton X-100 for 15 min at room temperature. The coverslips were incubated with anti-c-Myc mouse MAb and anti-HA rabbit PAb. The samples were washed three times with the wash buffer provided with the kit. The PLA probes (anti-mouse MINUS and anti-rabbit PLUS) were diluted in the antibody diluent provided with the kit, and the samples were incubated for 1 h at 37°C in a humidity chamber. To assess the interaction between HUWE1 and Nrf2, HepG2 cells seeded on glass coverslips placed in a 24-well plate were co-transfected with pCAG-HA-HUWE1 and pCAG-FLAG-Nrf2, and either pEGFP-C3 or pEGFP-C3-HBx. At 48 h posttransfection, the cells were fixed and permeabilized as described above. The coverslips were incubated with anti-FLAG M2 mouse MAb and anti-HA rabbit PAb. The PLA probes (anti-mouse MINUS and anti-rabbit PLUS) were applied for 1 h at 37°C. The samples were washed and processed according to the manufacturer’s instructions for probe ligation, signal amplification, and mounting. The samples were examined using a confocal laser scanning microscope (LSM 700; Carl Zeiss).

### Cell viability assay

Cell viability was evaluated by measuring intracellular ATP levels using the CellTiter-Glo v2.0 assay kit (Promega) according to the manufacturer’s protocol. Luminescence signal was measured using a GloMax Navigator microplate luminometer GM2000 (Promega).

### Statistical analysis

All quantitative data are presented as means ± standard errors of the mean (SEM) of at least three independent experiments. Statistical significance was determined using Student’s *t*-test, with *P* < 0.05 considered statistically significant.

## RESULTS

### HUWE1 participates in HBx-mediated K6-linked polyubiquitylation of Nrf2

To investigate whether HUWE1 participates in K6-linked polyubiquitylation of Nrf2, we performed cell-based ubiquitylation assays in HepG2 cells. Cells were co-transfected with pCAG-FLAG-Nrf2, and the plasmid pRK5-HA-Ubiquitin (Ub) encoding HA-tagged wild-type (WT)-Ub or pRK5-HA-Ub-K6, in which only ubiquitin lysine-6 is retained (K6), and either wild-type HBV or HBx-deficient HBV plasmid, pUC19-HBV-C-AT_JPN or pUC19-HBV-C-AT_JPN(ΔHBx), respectively (Fig. 1A). Immunoprecipitation with anti-FLAG antibody followed by immunoblotting with anti-HA antibody revealed that K6-linked polyubiquitylation of Nrf2 was markedly enhanced in the presence of HBx (Fig. 1A, top panel; compare lane 5 with lane 4). Notably, K6-linked Nrf2 polyubiquitylation was markedly lower in HBx-deficient HBV-replicating cells (ΔHBx) than in wild-type HBV-replicating cells (Fig. 1A, top panel; compare lane 6 with lane 5). These results suggest that HBx is required for K6-linked polyubiquitylation of Nrf2.

**Fig. 1.**
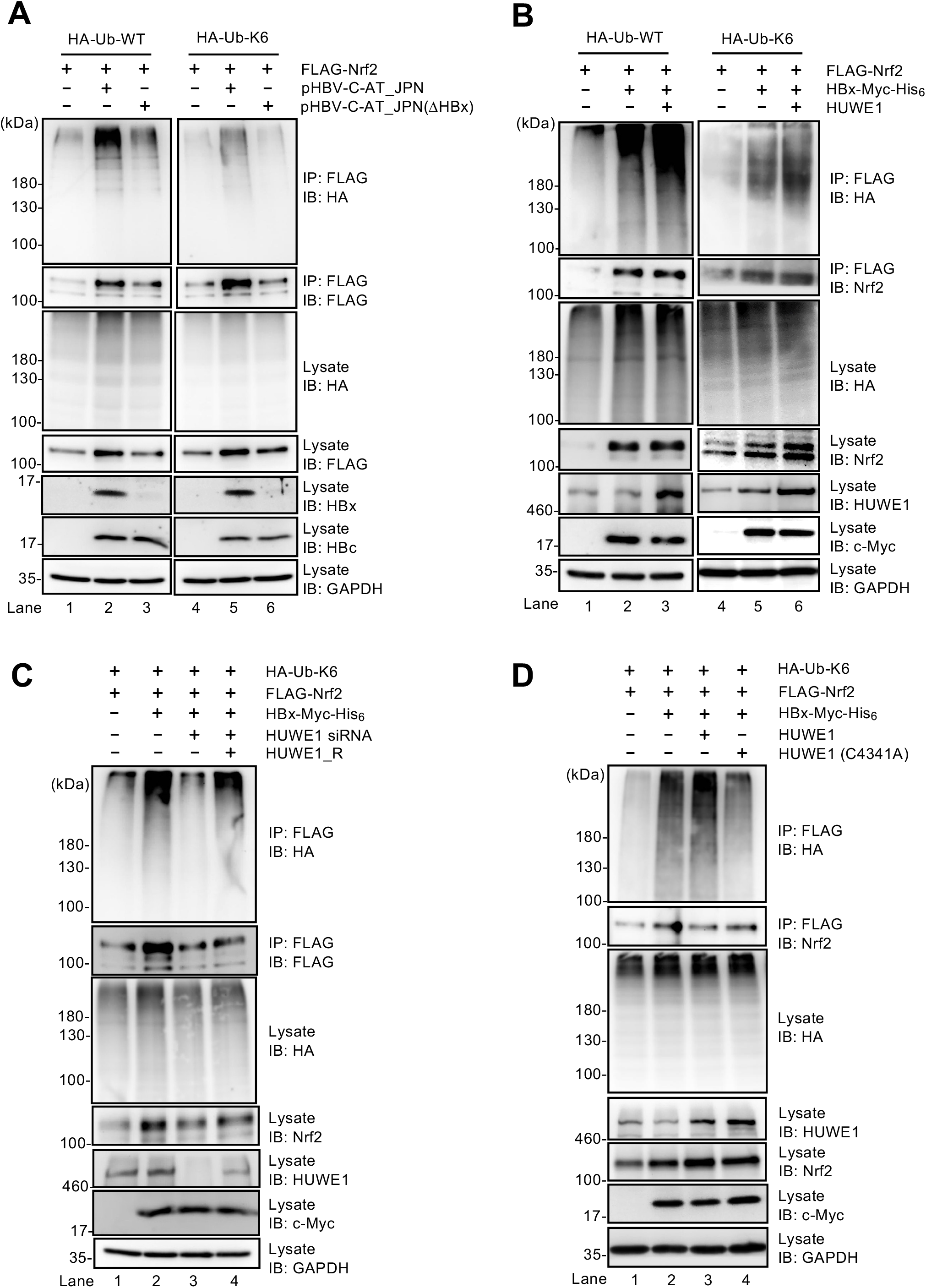
HUWE1 participates in HBx-mediated K6-linked polyubiquitylation of Nrf2. (A) HepG2 cells were co-transfected with pCAG-FLAG-Nrf2 and either pRK5-HA-Ub or pRK5-HA-Ub-K6 and with either pUC19-HBV-C-AT_JPN, carrying a 1.3-mer overlength HBV, or pUC19-HBV-C-AT_JPN(ΔHBx), carrying a 1.3-mer overlength HBV lacking HBx gene expression. At 48 h posttransfection, cells were harvested and lysed under denaturing conditions. Cell lysates were immunoprecipitated with anti-FLAG M2 affinity beads, followed by immunoblotting with anti-HA PAb (top panel) or anti-FLAG MAb (second panel). Input samples were immunoblotted with anti-HA PAb (third panel), anti-FLAG MAb (fourth panel), anti-HBx PAb (fifth panel), anti-HBc MAb (sixth panel), and anti-GAPDH MAb (seventh panel). GAPDH served as a loading control. (B) HepG2 cells were co-transfected with pCAG-FLAG-Nrf2, pEF1A-HBx-Myc-His_6_, pCAG-HUWE1, and either pRK5-HA-Ub or pRK5-HA-Ub-K6. At 48 h posttransfection, cells were harvested. Cell lysates were subjected to immunoprecipitation with anti-FLAG M2 affinity beads, followed by immunoblotting with anti-HA PAb (top panel) or anti-Nrf2 MAb (second panel). Input samples were immunoblotted with anti-HA PAb (third panel), anti-Nrf2 MAb (fourth panel), anti-HUWE1 PAb (fifth panel), anti-c-Myc MAb (sixth panel), and anti-GAPDH MAb (seventh panel). GAPDH served as a loading control. (C) HepG2 cells were transfected with 40 nM HUWE1 siRNA or control siRNA. At 24 h post-siRNA transfection, cells were co-transfected with pCAG-FLAG-Nrf2, pRK5-HA-Ub-K6, pEF1A-HBx-Myc-His_6_, and the HUWE1 siRNA-resistant plasmid pCAG-HUWE1_R. At 48 h posttransfection, cells were harvested. Cell lysates were subjected to immunoprecipitation with anti-FLAG M2 affinity beads, followed by immunoblotting with anti-HA PAb (top panel) or anti-FLAG MAb (second panel). Input samples were immunoblotted with anti-HA PAb (third panel), anti-Nrf2 MAb (fourth panel), anti-HUWE1 PAb (fifth panel), anti-c-Myc MAb (sixth panel), or anti-GAPDH MAb (seventh panel). GAPDH served as a loading control. HUWE1_R, siRNA-resistant HUWE1. (D) HepG2 cells were co-transfected with pCAG-FLAG-Nrf2, pRK5-HA-Ub-K6, pEF1A-HBx-Myc-His_6_ and either pCAG-HUWE1(WT) or the catalytically inactive mutant pCAG-HUWE1(C4341A). At 48 h posttransfection, cells were harvested. Cell lysates were subjected to immunoprecipitation with anti-FLAG M2 affinity beads, followed by immunoblotting with anti-HA PAb (top panel) or anti-Nrf2 MAb (second panel). Input samples were immunoblotted with anti-HA PAb (third panel), anti-HUWE1 PAb (fourth panel), anti-Nrf2 MAb (fifth panel), anti-c-Myc MAb (sixth panel), or anti-GAPDH MAb (seventh panel). GAPDH served as a loading control. Data in panels A to D are representative of three independent experiments.

To determine whether HUWE1 contributes to this process, HepG2 cells were co-transfected with pCAG-HUWE1, pCAG-FLAG-Nrf2, pEF1A-HBx-Myc-His_6_, and either pRK5-HA-Ub or pRK5-HA-Ub-K6. Overexpression of wild-type HUWE1 further enhanced HBx-dependent K6-linked polyubiquitylation of Nrf2 (Fig. 1B, top panel; compare lane 6 with lane 5). Conversely, knockdown of endogenous HUWE1 using siRNA substantially reduced K6-linked polyubiquitylation of Nrf2 (Fig. 1C, top panel; compare lane 3 with lane 2), whereas re-expression of an siRNA-resistant HUWE1 (HUWE1_R) plasmid restored it (Fig. 1C, top panel, lane 4), indicating the specificity of the siRNA effect. Importantly, overexpression of the catalytically inactive HUWE1 (C4341A) mutant failed to enhance K6-linked polyubiquitylation of Nrf2 (Fig. 1D, top panel, lane 4), indicating that the HUWE1 E3 ligase activity is required for K6-linked polyubiquitylation of Nrf2.

### HUWE1 specifically interacts with HBx

To examine whether HUWE1 physically interacts with HBx, we performed coimmunoprecipitation analysis. Co-transfection of pCAG-HA-HUWE1 and pEF1-HBx-Myc-His_6_ into HepG2 cells, followed by immunoprecipitation with anti-HA antibody and immunoblot analysis with anti-c-Myc antibody demonstrated a specific interaction between HUWE1 and HBx (Fig. 2A, top panel, lane 4). In contrast, coimmunoprecipitation analysis of cells co-expressing HA-HUWE1 and HBc-Myc-His_6_ showed that HUWE1 did not interact with HBc (Fig. 2B, top panel, lane 3), supporting the specificity of the HUWE1–HBx interaction.

**Fig. 2.**
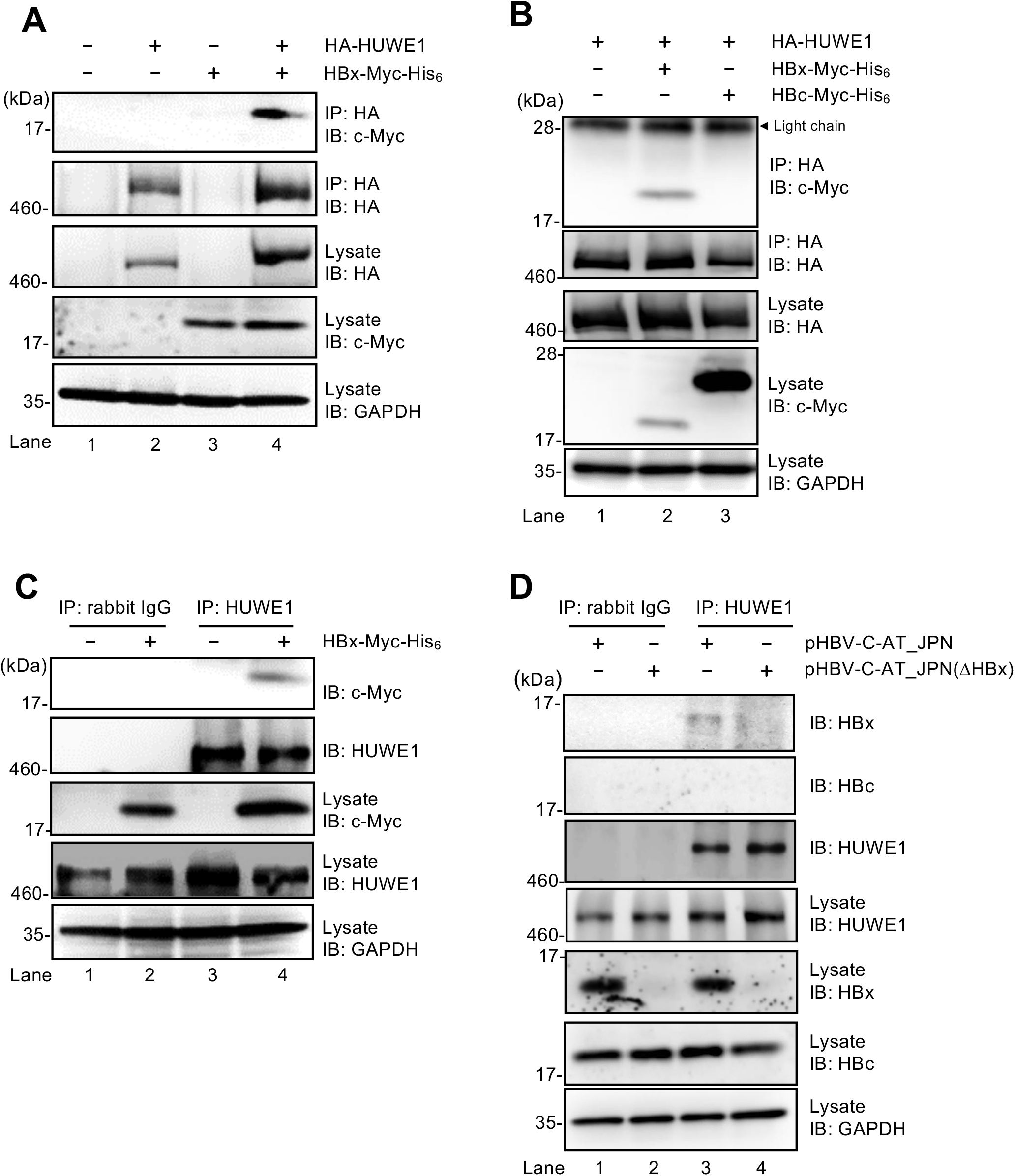
HUWE1 interacts with HBx. (A) HepG2 cells were co-transfected with pCAG-HA-HUWE1 and pEF1A-HBx-Myc-His_6_. At 48 h posttransfection, cells were harvested. Cell lysates were immunoprecipitated with anti-HA PAb, followed by immunoblotting with anti-c-Myc MAb (top panel) or anti-HA PAb (second panel). Input samples were immunoblotted with anti-HA PAb (third panel), anti-c-Myc MAb (fourth panel), and anti-GAPDH MAb (fifth panel). GAPDH served as a loading control. (B) HepG2 cells were co-transfected with pCAG-HA-HUWE1 and either pEF1A-HBx-Myc-His_6_ or pEF1A-HBc-Myc-His_6_. At 48 h posttransfection, cells were harvested. Cell lysates were immunoprecipitated with anti-HA PAb, followed by immunoblotting with anti-c-Myc MAb (top panel) or anti-HA PAb (second panel). Input samples were immunoblotted with anti-HA PAb (third panel), anti-c-Myc MAb (fourth panel), and anti-GAPDH MAb (fifth panel). GAPDH served as a loading control. (C) HepG2 cells were transfected with pEF1A-HBx-Myc-His_6_. At 48 h posttransfection, cells were harvested. Cell lysates were immunoprecipitated with anti-HUWE1 PAb or rabbit IgG (control), followed by immunoblotting with anti-c-Myc MAb (top panel) or anti-HUWE1 PAb (second panel). Input samples were immunoblotted with anti-c-Myc MAb (third panel), anti-HUWE1 PAb (fourth panel), and anti-GAPDH MAb (fifth panel). GAPDH served as a loading control. (D) HepG2 cells were transfected with pUC19-HBV-C-AT_JPN or pUC19-HBV-C-AT_JPN(ΔHBx). At 48 h posttransfection, cells were harvested. Cell lysates were immunoprecipitated with anti-HUWE1 PAb or rabbit IgG (control), followed by immunoblotting with anti-HBx PAb (top panel), anti-HBc MAb (second panel), or anti-HUWE1 PAb (third panel). Input samples were immunoblotted with anti-HUWE1 PAb (fourth panel), anti-HBx PAb (fifth panel), anti-HBc MAb (sixth panel), and anti-GAPDH MAb (seventh panel). GAPDH served as a loading control. Data in panels A to D are representative of three independent experiments.

To investigate the interaction between endogenous HUWE1 and HBx, HepG2 cells were transfected with pEF1A-HBx-Myc-His_6_. Consistent with the results obtained in the HUWE1 overexpression experiments, immunoprecipitation analysis using an anti-HUWE1 antibody demonstrated that HBx-Myc-His_6_ was coimmunoprecipitated with endogenous HUWE1 (Fig. 2C, top panel, lane 4), whereas a non-specific IgG control demonstrated no signal (Fig. 2C, top panel, lane 2).

To further verify the interaction between endogenous HUWE1 and HBx in HBV-replicating cells, HepG2 cells were transfected with pUC19-HBV-C-AT_JPN or pUC19-HBV-C-AT_JPN(ΔHBx). Immunoprecipitation analysis using anti-HUWE1 antibody demonstrated that HBx (Fig. 2D, top panel, lane 3), but not HBc (Fig. 2D, second panel, lane 3), was coimmunoprecipitated with HUWE1. No band was detected in HBV-C-AT_JPN(ΔHBx)-replicating cells (Fig. 2D, top panel, lane 4), confirming the specificity of HUWE1 binding to HBx. These results indicate that HUWE1 specifically interacts with HBx.

### HUWE1 colocalizes with HBx predominantly in the cytoplasm

To determine the subcellular localization of the HUWE1−HBx interaction, we performed indirect immunofluorescence staining and in situ proximity ligation assay (PLA). Immunofluorescence staining of the cells expressing HA-HUWE1 (green) and HBx-Myc-His_6_ (red) revealed co-localization predominantly in the cytoplasm, with minimal nuclear signal (Fig. 3A, bottom panel, merge). Consistent with this, PLA using anti-HA and anti-c-Myc antibodies produced cytoplasmic PLA signals (red dots) that were absent in single-protein controls (Fig. 3B). These results suggest that HUWE1 interacts with HBx predominantly in the cytoplasm.

**Fig. 3.**
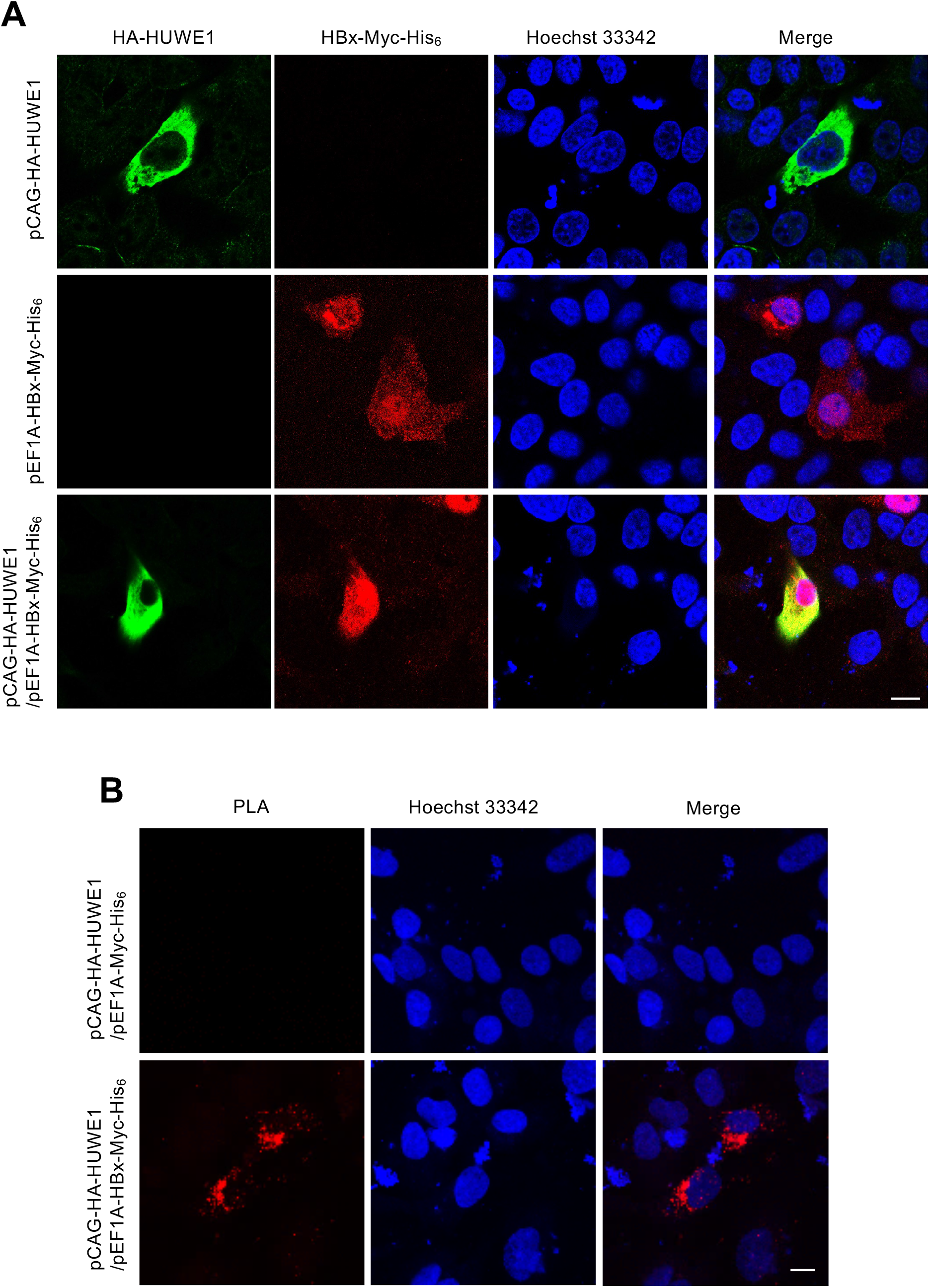
HUWE1 colocalizes with HBx predominantly in the cytoplasm. (A) HepG2 cells were transfected with pCAG-HA-HUWE1 alone, pEF1A-HBx-Myc-His_6_ alone, or both plasmids. At 48 h posttransfection, cells were fixed and subjected to indirect immunofluorescence staining with anti-HUWE1 rabbit PAb followed by Alexa Fluor 488-conjugated goat anti-rabbit IgG (green), and anti-c-Myc mouse MAb followed by Alexa Fluor 594-conjugated goat anti-mouse IgG (red). Nuclei were counterstained with Hoechst 33342 (blue). Scale bar, 10 μm. (B) HepG2 cells were co-transfected with pCAG-HA-HUWE1 and either pEF1A-Myc-His_6_ (upper panel) or pEF1A-HBx-Myc-His_6_ (lower panel). At 48 h posttransfection, cells were fixed and subjected to an in situ proximity ligation assay (PLA) using both anti-HA rabbit PAb and anti-c-Myc mouse MAb. Nuclei were counterstained with Hoechst 33342 (blue). Scale bar, 10 μm. Data in panels A and B are representative of three independent experiments.

### HUWE1-mediated K6-linked polyubiquitylation of Nrf2 depends on the interaction between HUWE1 and HBx

To determine whether HUWE1 binds Nrf2 directly or requires HBx as a scaffold, we performed coimmunoprecipitation experiments by co-expressing HA-HUWE1, FLAG-Nrf2, and HBx-Myc-His_6_ in HepG2 cells. Immunoprecipitation with anti-HA antibody pulled down both FLAG-Nrf2 and HBx-Myc-His_6_ (Fig. 4A, first and second panels, lane 3), suggesting the formation of a ternary HUWE1/HBx/Nrf2 complex.

**Fig. 4.**
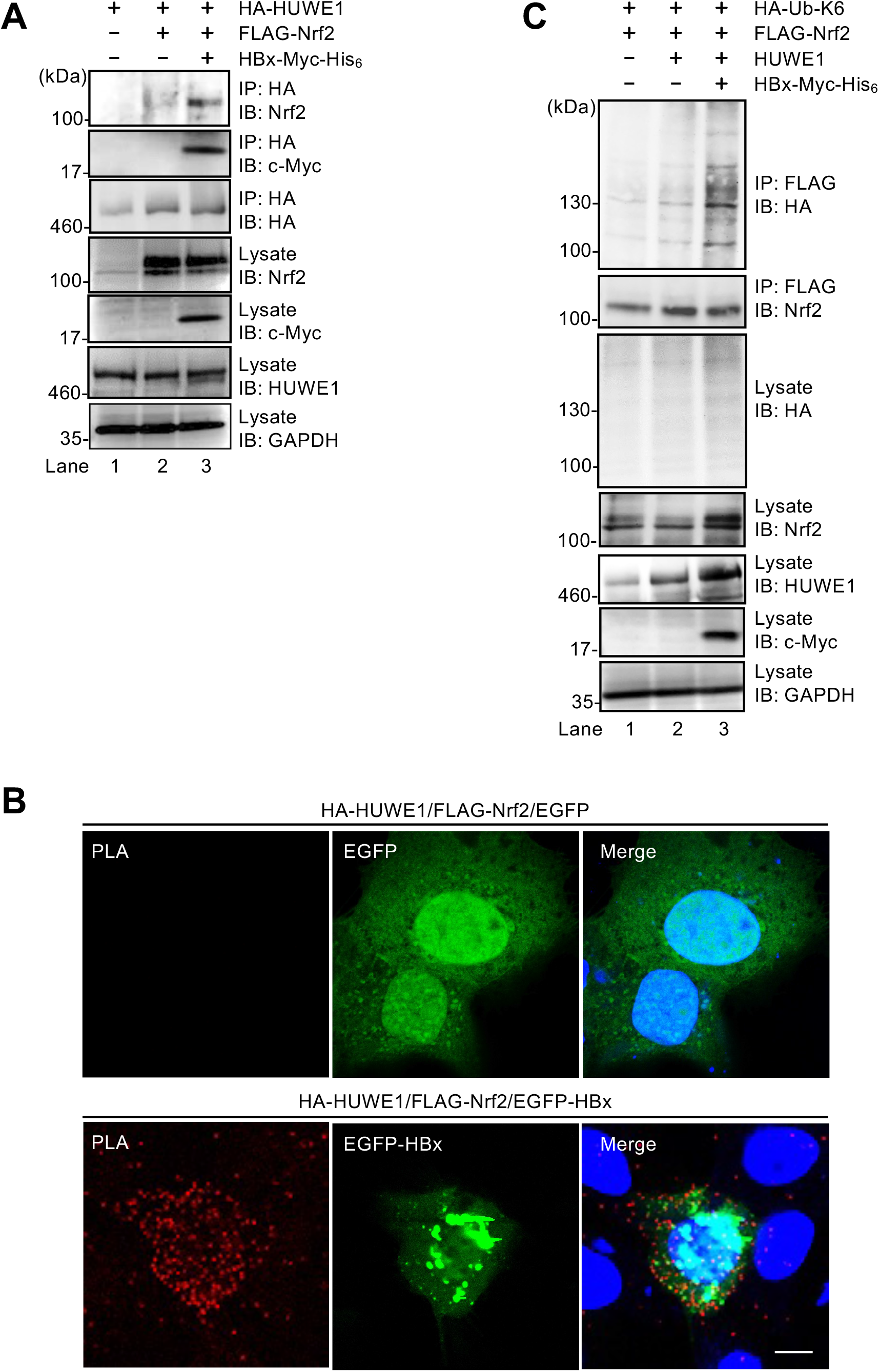
HUWE1-mediated K6-linked polyubiquitylation of Nrf2 depends on the interaction between HUWE1 and HBx. (A) HepG2 cells were co-transfected with pCAG-HA-HUWE1, pCAG-FLAG-Nrf2, and pEF1A-HBx-Myc-His_6_. At 48 h posttransfection, cells were harvested. Cell lysates were immunoprecipitated with anti-HA PAb, followed by immunoblotting with anti-Nrf2 MAb (top panel), anti-c-Myc MAb (second panel), and anti-HUWE1 PAb (third panel). Input samples were immunoblotted with the indicated antibodies, and GAPDH served as a loading control. (B) HepG2 cells were co-transfected with pCAG-FLAG-Nrf2 and pCAG-HA-HUWE1 and either pEGFP-C3 (upper panel) or pEGFP-C3-HBx (lower panel). At 48 h posttransfection, the cells were fixed and incubated with anti-FLAG mouse MAb and anti-HA rabbit PAb, followed by an in situ PLA. Scale bar, 10 μm. (C) HepG2 cells were co-transfected with pCAG-FLAG-Nrf2 and pRK5-HA-Ub-K6, together with pCAG-HUWE1 and/or pEF1A-HBx-Myc-His_6_ as indicated. At 48 h posttransfection, cells were harvested. Cell lysates were immunoprecipitated with anti-FLAG M2 affinity beads, followed by immunoblotting with anti-HA PAb (top panel) or anti-Nrf2 MAb (second panel). Input samples were immunoblotted with anti-HA PAb (third panel), anti-Nrf2 MAb (fourth panel), anti-HUWE1 PAb (fifth panel), anti-c-Myc MAb (sixth panel), or anti-GAPDH MAb (seventh panel). GAPDH served as a loading control. Data in panels A to C are representative of three independent experiments.

To further determine whether the HUWE1-Nrf2 interaction depends on HBx, we performed in situ PLA analyses. HepG2 cells were co-transfected with pCAG-HA-HUWE1, pCAG-FLAG-Nrf2, and either pEGFP-C3 or pEGFP-C3-HBx. PLA signals between HA-HUWE1 and FLAG-Nrf2 were detected only in the presence of HBx (Fig. 4B, lower panel), suggesting that HBx facilitates the association between HUWE1 and Nrf2.

To determine whether HBx is required for HUWE1-catalyzed K6-linked polyubiquitylation of Nrf2, we performed cell-based ubiquitylation assays in the presence or absence of HBx expression. HepG2 cells were co-transfected with pCAG-FLAG-Nrf2, pRK5-HA-Ub-K6, pCAG-HUWE1, and pEF1A-HBx-Myc-His_6_. Cell lysates were immunoprecipitated with anti-FLAG beads and immunoblotted with an anti-HA polyclonal antibody to detect ubiquitylated FLAG-Nrf2. Immunoblot analysis revealed that overexpression of HUWE1 alone had little effect on K6-linked polyubiquitylation of Nrf2 (Fig. 4C, top panel, lane 2). However, the level of K6-linked polyubiquitylation of Nrf2 was increased when HUWE1 was co-expressed with HBx-Myc-His_6_ (Fig. 4C, top panel, lane 3). These results suggest that HBx is required for HUWE1-mediated K6-linked polyubiquitylation of Nrf2.

### HUWE1 contributes to the stabilization of Nrf2 in an HBx-dependent manner

To investigate whether HUWE1-mediated K6-linked polyubiquitylation of Nrf2 affects Nrf2 stability, we performed cycloheximide (CHX) chase analyses. HepG2 cells were co-transfected with pCAG-FLAG-Nrf2 and either pEF1A-Myc-His_6_ (Fig. 5A and B) or pEF1A-HBx-Myc-His_6_ (Fig. 5C and D) in the presence or absence of HUWE1 siRNA. Cells were collected at 0, 1, 2, 4, and 6 h after treatment with 100 μg/mL CHX and analyzed by immunoblotting. CHX chase analysis demonstrated that HUWE1 knockdown had no significant effect on Nrf2 stability in cells lacking HBx (Fig. 5A, top panel, and Fig. 5B). In contrast, in HBx-expressing cells, HUWE1 knockdown markedly accelerated Nrf2 degradation compared with that in the control siRNA-transfected cells (Fig. 5C, top panel, and Fig. 5D), suggesting that HUWE1-mediated K6-linked polyubiquitylation stabilizes Nrf2 specifically in the presence of HBx, which is consistent with the notion that HUWE1 mediates Nrf2 stabilization in an HBx-dependent manner.

**Fig. 5.**
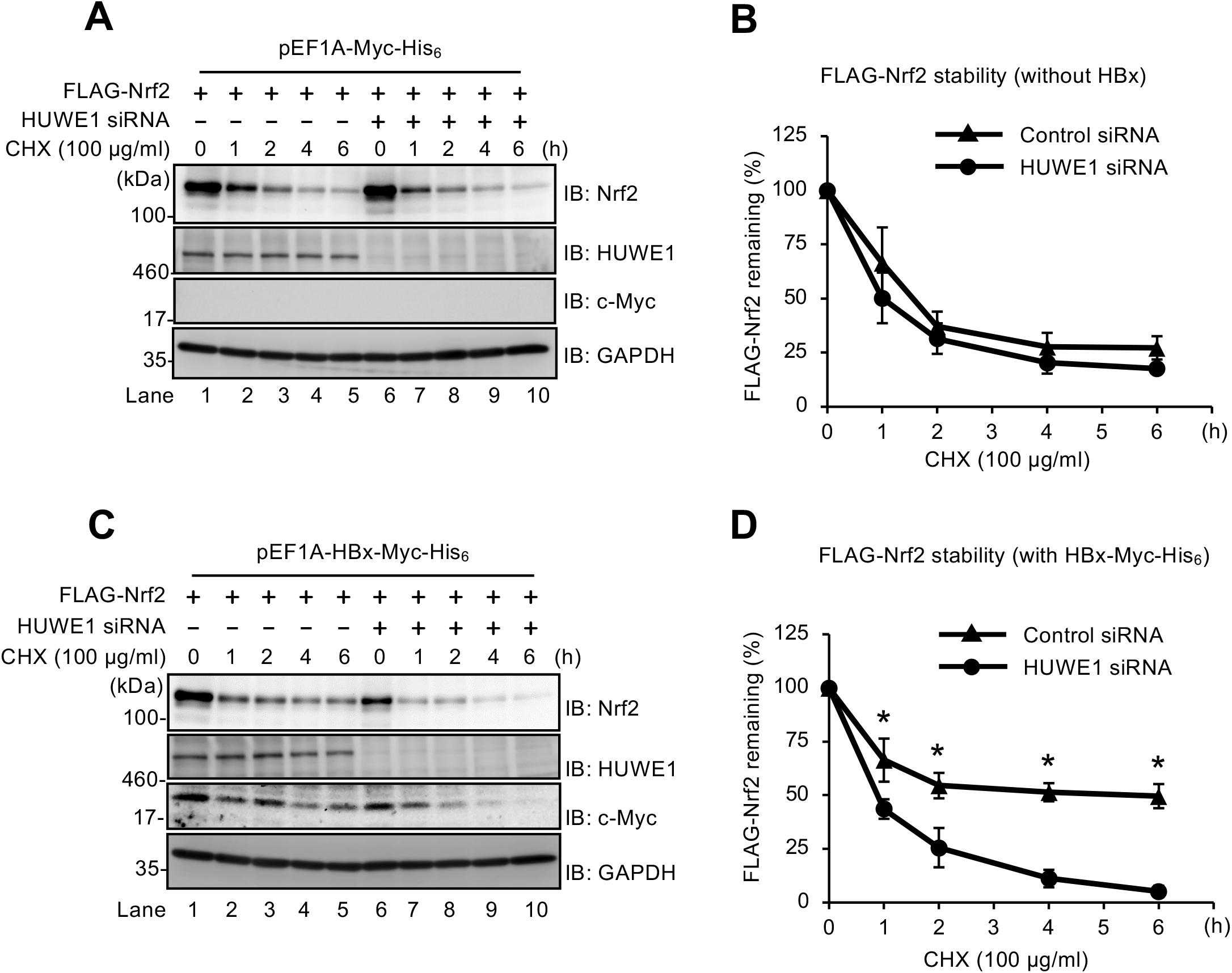
HUWE1 contributes to the stabilization of Nrf2 in an HBx-dependent manner. (A to D) HepG2 cells were transfected with 40 nM HUWE1 siRNA or control siRNA. At 24 h post-siRNA transfection, cells were co-transfected with pCAG-FLAG-Nrf2 and either pEF1A-Myc-His_6_ (panels A and B) or pEF1A-HBx-Myc-His_6_ (panels C and D). At 48 h posttransfection, cells were treated with 100 µg/mL cycloheximide (CHX) for 0, 1, 2, 4, or 6 h prior to harvesting. Cell lysates were then subjected to immunoblotting with the indicated antibodies. GAPDH served as a loading control. (B, D) Specific signals were quantified by densitometry. The quantifications were performed using ImageJ, normalized to the corresponding loading controls. The percentage of remaining FLAG-Nrf2 at each time point was compared with that at time zero. Closed triangles (▴) indicate control siRNA, and closed circles (●) indicate HUWE1 siRNA. Data are mean ± SEM of three independent experiments. \**P* < 0.05 compared with the control siRNA by Student’s t test.

### Knockdown of HUWE1 enhances HBV replication

To examine the functional significance of HUWE1 in HBV replication, we performed HUWE1 knockdown experiments in multiple HBV replication models. In HepG2 cells transfected with pUC19-HBV-D-IND60, carrying a 1.3-mer overlength HBV, HUWE1 knockdown significantly increased both HBV RNA (Fig. 6A) and pregenomic RNA (pgRNA) (Fig. 6B) levels compared with control siRNA. Re-expression of siRNA-resistant HUWE1 reduced HBV RNA levels (Fig. 6A) and pgRNA levels (Fig. 6B), confirming on-target specificity. Immunoblot analysis confirmed effective HUWE1 knockdown (Fig. 6C, top panel, lane 2) and the corresponding increase in HBc protein level (Fig. 6C, middle panel, lane 2).

**Fig. 6.**
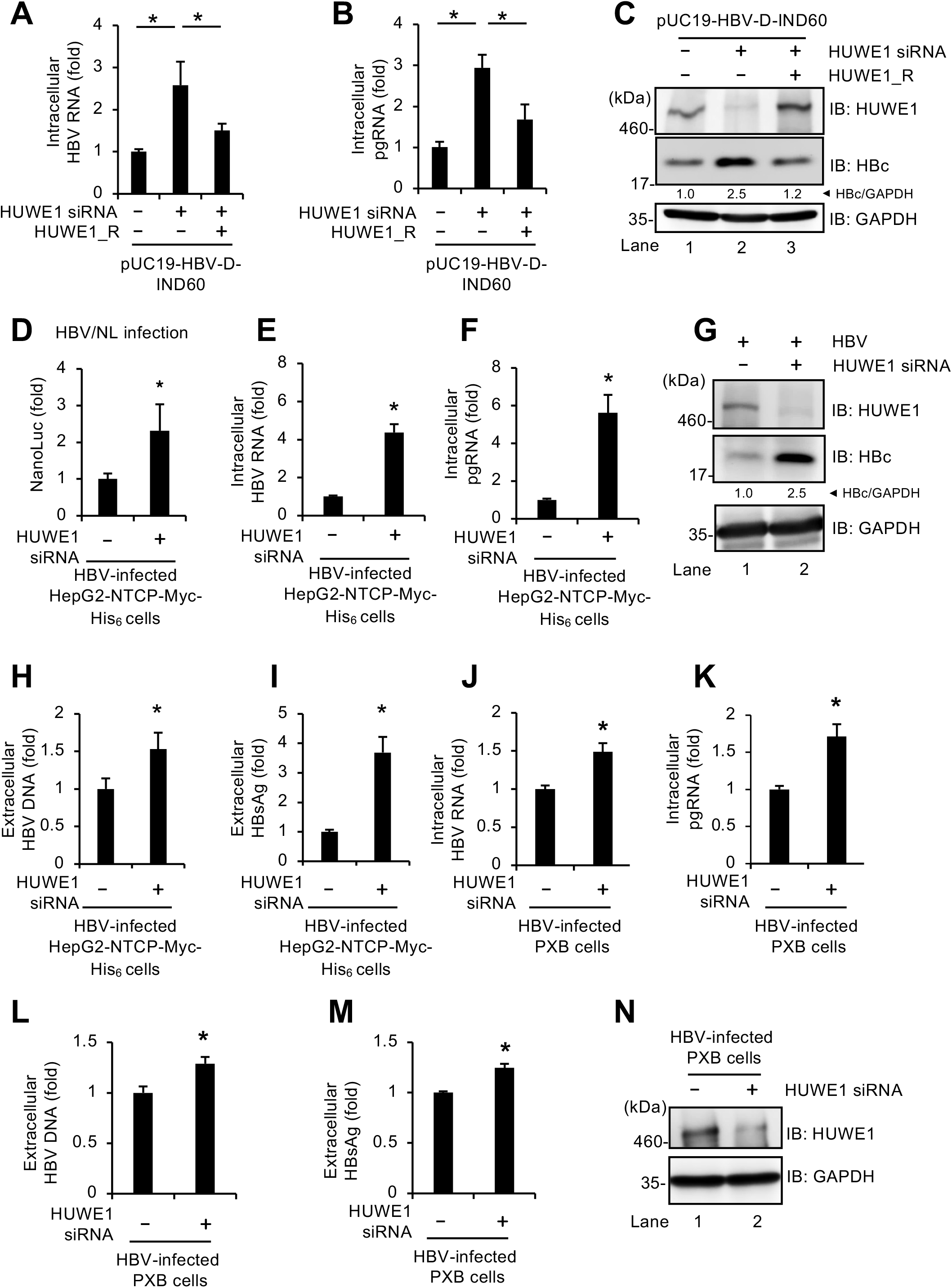
Knockdown of HUWE1 enhances HBV replication. (A to C) HepG2 cells were transfected with 40 nM HUWE1 siRNA or control siRNA. At 24 h post-siRNA transfection, cells were co-transfected with pUC19-HBV-D-IND60 and the HUWE1 siRNA-resistant plasmid pCAG-HUWE1_R. At 48 h post-plasmid transfection, cells were harvested. Total RNA was extracted, and HBV RNA (A) and pgRNA (B) were quantified by RT-qPCR and normalized to GAPDH mRNA. Data represent the means ± SEM from three biological replicates. Values for control cells were arbitrarily set to 1.0. \**P* < 0.05 by Student’s t test. HUWE1_R, siRNA-resistant HUWE1. (C) Cell lysates were subjected to immunoblotting using anti-HUWE1 PAb (top panel), anti-HBc MAb (middle panel), and anti-GAPDH MAb (bottom panel), with GAPDH as the loading control. The relative levels of HBc were quantified by densitometry and are indicated below each lane. (D) HepG2-NTCP-Myc-His_6_ cells were transfected with 40 nM HUWE1 siRNA or control siRNA. At 48 h posttransfection, cells were infected with recombinant HBV expressing the NanoLuc (NL) reporter gene (HBV/NL) at 100 genome equivalents (GEq)/cell. At 4 days postinfection (dpi), cells were lysed to measure NL activity as an indicator of HBV replication. Results represent the mean ± SEM from three biological replicates. Values for control cells were arbitrarily set to 1.0. \**P* < 0.05 by Student’s t test. (E-I) HepG2-NTCP-Myc-His_6_ cells were transfected with 40 nM HUWE1 siRNA or control siRNA. At 48 h posttransfection, cells were infected with HBV at 1,000 GEq/cell, and harvested at 5 dpi. Total RNA was extracted, and HBV RNA (E) and pgRNA (F) were quantified by RT-qPCR. Data represent the means ± SEM from three biological replicates. Values for control cells were arbitrarily set to 1.0. \**P* < 0.05 by Student’s t test. (G) Cell lysates were subjected to immunoblotting using anti-HUWE1 PAb (top panel), anti-HBc MAb (middle panel), and anti-GAPDH MAb (bottom panel), with GAPDH as the loading control. The relative levels of HBc were quantified by densitometry and are indicated below each lane. Extracellular HBV DNA (H) and HBsAg (I) levels were quantified by qPCR and ELISA, respectively. Data represent the means ± SEM from three biological replicates. Values for control cells were arbitrarily set to 1.0. \**P* < 0.05 by Student’s t test. (J-N) PXB cells were transfected with 40 nM HUWE1 siRNA or control siRNA. At 48 h posttransfection, cells were infected with HBV at 1,000 GEq/cell. At 24 h postinfection and 5 dpi, the cells were re-transfected with siRNA and maintained until harvesting. At 8 dpi, total RNA was extracted, and HBV RNA (J) and pgRNA (K) were quantified by RT-qPCR. Data represent the means ± SEM from three biological replicates. Values for control cells were arbitrarily set to 1.0. \**P* < 0.05 by Student’s t test. Extracellular HBV DNA (L) and HBsAg (M) levels were quantified by qPCR and ELISA, respectively. Data represent the means ± SEM from three biological replicates. Values for control cells were arbitrarily set to 1.0. \**P* < 0.05 by Student’s t test. (N) The expression of HUWE1 in cell lysates was analyzed by immunoblotting with anti-HUWE1 PAb (top panel). GAPDH was detected using anti-GAPDH MAb (bottom panel) as the loading control.

To further examine the role of HUWE1 in HBV replication, we used recombinant HBV particles carrying a chimeric HBV genome encoding NanoLuc (NL), known as the HBV/NL system. HepG2-NTCP-Myc-His_6_ cells were infected with HBV/NL in the presence or absence of HUWE1 siRNA. At 4 dpi, cells were lysed and NanoLuc activity was measured. The HBV/NL assay demonstrated that knockdown of HUWE1 significantly increased intracellular NL activity (Fig. 6D).

Next, we assessed the role of HUWE1 in HBV replication in HBV-infected HepG2-NTCP-Myc-His_6_ cells. Cells were infected with HBV in the presence or absence of HUWE1 siRNA. The RT-qPCR analysis demonstrated that HUWE1 knockdown significantly enhanced the levels of intracellular HBV RNA (Fig. 6E) and pgRNA (Fig. 6F). The level of HBc protein was consistently increased by knockdown of HUWE1 (Fig. 6G, middle panel, lane 2). Moreover, extracellular levels of HBV DNA (Fig. 6H) and HBsAg (Fig. 6I) were significantly increased following HUWE1 knockdown. Importantly, HUWE1 knockdown also enhanced intracellular levels of HBV RNA (Fig. 6J) and pgRNA (Fig. 6K), as well as extracellular levels of HBV DNA (Fig. 6L) and HBsAg (Fig. 6M), in HBV-infected PXB cells, primary human hepatocytes derived from humanized chimeric mouse livers. We confirmed that endogenous HUWE1 was efficiently knocked down by siRNA (Fig. 6N, top panel, lane 2). Collectively, these results suggest that HUWE1 contributes to the suppression of HBV replication in these experimental models.

### The HUWE1 E3 ligase inhibitor BI8626 suppresses K6-linked polyubiquitylation of Nrf2 and enhances HBV replication

To further evaluate the functional contribution of HUWE1, we examined the effects of the HUWE1 E3 ligase inhibitor BI8626 on the K6-linked polyubiquitylation of Nrf2 and HBV replication. HepG2 cells were co-transfected with pCAG-FLAG-Nrf2, pRK5-HA-Ub-K6, and either pEF1A-HBx-Myc-His_6_ or pEF1A-Myc-His_6_. Treatment of HepG2 cells with 10 µM BI8626 clearly reduced HBx-dependent K6-linked polyubiquitylation of Nrf2 (Fig. 7A, top panel, compare lane 4 with lane 3), without affecting cell viability at this concentration (Fig. 7B).

**Fig. 7.**
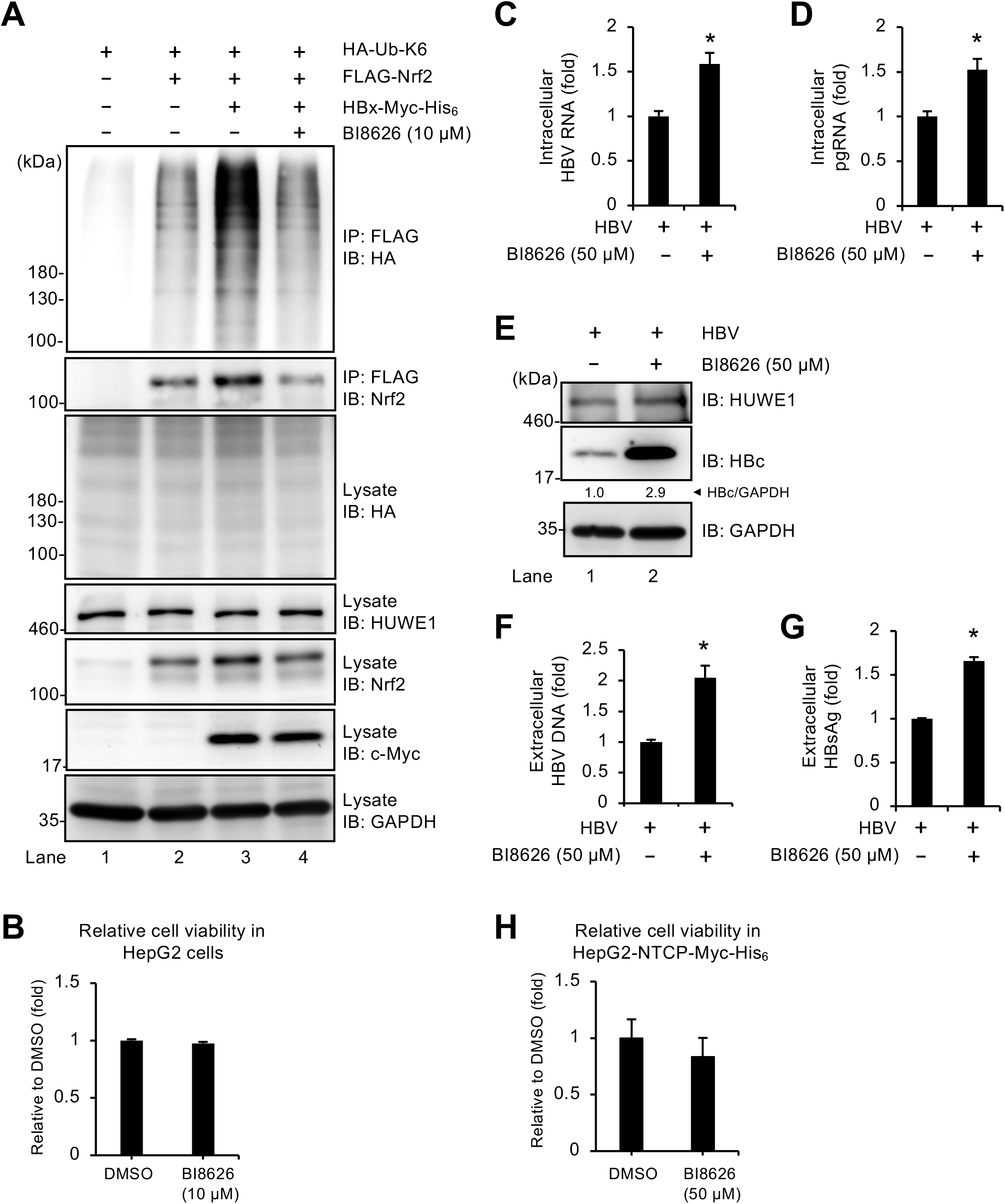
HUWE1 E3 ligase inhibitor BI8626 enhances HBV replication by suppressing HUWE1-mediated K6-linked polyubiquitylation of Nrf2. (A) HepG2 cells were co-transfected with pCAG-FLAG-Nrf2, pRK5-HA-Ub-K6, and pEF1A-HBx-Myc-His_6_. At 24 h posttransfection, cells were treated with 10 µM BI8626 or equivalent volume of dimethyl sulfoxide (DMSO; vehicle control) for 48 h prior to harvesting. Cell lysates were immunoprecipitated with anti-FLAG M2 affinity beads and subjected to immunoblotting with anti-HA PAb (top panel) or anti-Nrf2 MAb (second panel). Input samples were immunoblotted with the indicated antibodies. GAPDH served as a loading control. (B) Cell viability of HepG2 cells treated with 10 µM BI8626 for 48 h, assessed by CellTiter-Glo v2.0 ATP assay. Results represent the mean values ± SEM from three biological replicates. (C to H) HepG2-NTCP-Myc-His_6_ cells were infected with HBV at 1,000 GEq/cell. At 24 h postinfection, cells were treated with 50 µM BI8626 and maintained until harvesting. At 10 dpi, total RNA was extracted, and HBV RNA (C) and pgRNA (D) were quantified by RT-qPCR. Data represent the means ± SEM from three biological replicates. Values for control cells were arbitrarily set to 1.0. \**P* < 0.05 by Student’s t test. (E) Cell lysates were analyzed by immunoblotting with anti-HUWE1 PAb (top panel), anti-HBc MAb (middle panel), and anti-GAPDH MAb (bottom panel), with GAPDH serving as a loading control. The relative levels of HBc were quantified by densitometry and are indicated below each lane. Extracellular HBV DNA (F) and HBsAg (G) levels were quantified by qPCR and ELISA, respectively. Data represent the means ± SEM from three biological replicates. Values for control cells were arbitrarily set to 1.0. \**P* < 0.05 by Student’s t test. (H) Cell viability of HepG2-NTCP-Myc-His_6_ cells treated with 50 µM BI8626 for 10 days, assessed by CellTiter-Glo 2.0. Data represent the means ± SEM from three biological replicates.

To evaluate the effect of BI8626 on HBV replication, HBV-infected HepG2-NTCP-Myc-His_6_ cells were treated with 50 µM BI8626 from 1 dpi until 10 dpi. Cells were harvested at 10 dpi. Inhibition of HUWE1 E3 ligase activity by BI8626 significantly enhanced the levels of intracellular HBV RNA (Fig. 7C), pgRNA (Fig. 7D), and HBc protein (Fig. 7E, middle panel, lane 2), as well as extracellular HBV DNA (Fig. 7F), and HBsAg (Fig.7G) in HBV-infected cells, without affecting cell viability at this concentration (Fig. 7H). Collectively, these results indicate that pharmacological inhibition of HUWE1 enhanced HBV replication, consistent with a role for HUWE1 E3 ligase activity in limiting HBV replication.

## DISCUSSION

In this study, we demonstrated that the HECT-type E3 ubiquitin ligase HUWE1 mediates HBx-dependent K6-linked polyubiquitylation of Nrf2, leading to Nrf2 stabilization and suppression of HBV replication. Our findings identify HUWE1 as a candidate host restriction factor against HBV and reveal a non-canonical K6-ubiquitin switch as an antiviral mechanism (Fig. 8).

**Fig. 8.**
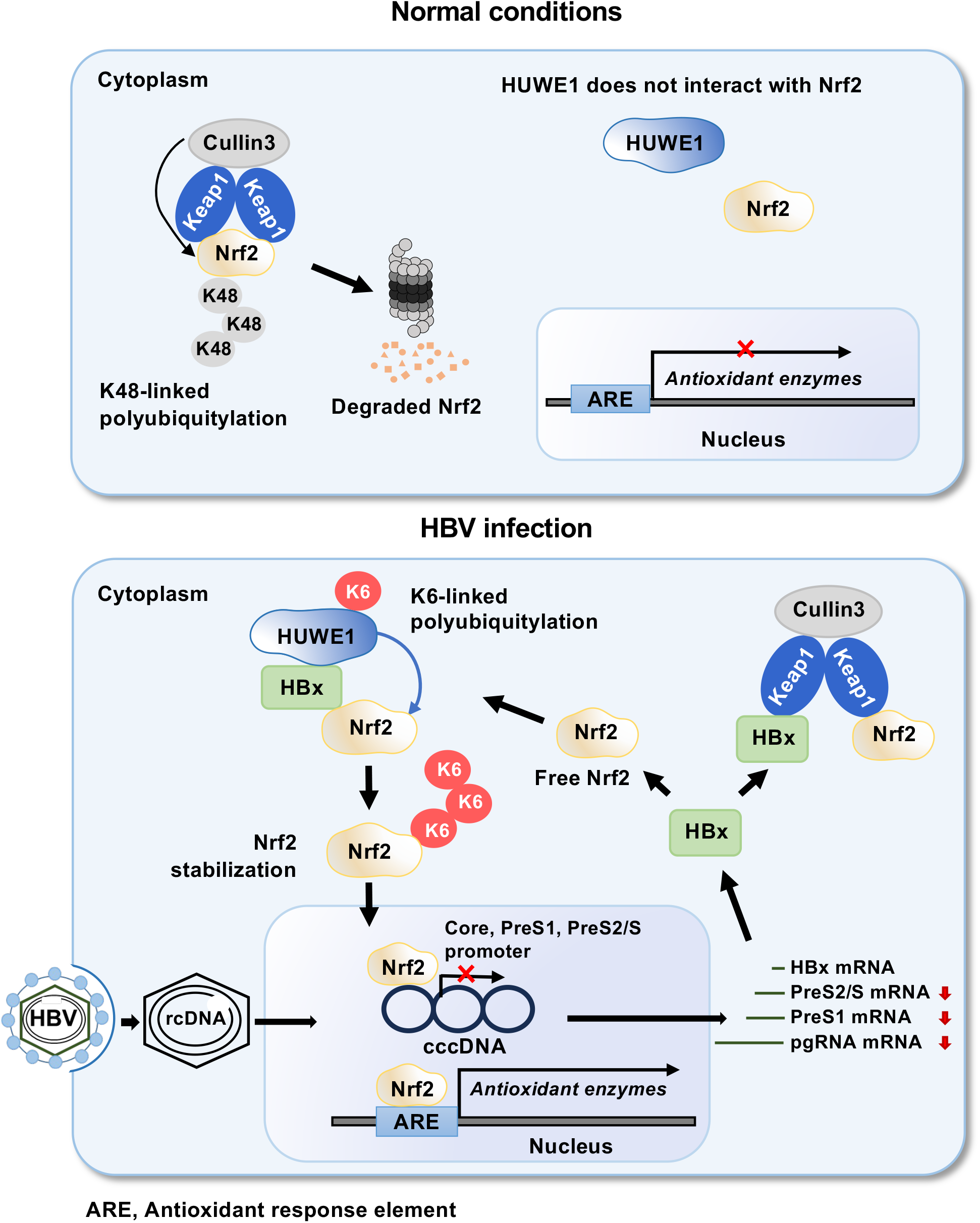
Proposed mechanism of HUWE1-mediated K6-linked polyubiquitylation of Nrf2 promoting its stabilization and suppression of HBV replication. Under normal conditions (upper panel), HUWE1 does not associate with Nrf2. Keap1 bridges Nrf2 to the Cullin3 E3 ligase complex, leading to K48-linked polyubiquitylation and proteasomal degradation of Nrf2. Upon HBV infection (lower panel), our working model proposes that HBx promotes the association of HUWE1 with Nrf2 in the cytoplasm. HUWE1 is proposed to enhance K6-linked polyubiquitylation of Nrf2, favoring a non-canonical modification state associated with Nrf2 stabilization. Stabilized Nrf2 may then translocate to the nucleus, activate antioxidant response element (ARE)-driven genes, and suppress HBV core, PreS1, and PreS2/S promoter activities, thereby contributing to suppression of HBV replication [11].

K6-linked polyubiquitylation has been shown to serve non-degradative functions, in contrast to the canonical K48-linked chains that target proteins for proteasomal degradation [19, 20, 35, 36]. K6-linked ubiquitin chains have been implicated in microtubule dynamics [37], mitophagy via Parkin [38], and innate immune signaling [39–42]. Our results add to this growing body of evidence by showing that HUWE1-mediated K6-linked polyubiquitylation stabilizes Nrf2 during HBV infection, representing a previously unrecognized antiviral function of K6-linked ubiquitin chains.

HUWE1 is a 482-kDa HECT domain E3 ubiquitin ligase known to catalyze K48-, K63-, and K6-linked polyubiquitylation [17, 43]. The HECT domain of HUWE1 forms a thioester intermediate with ubiquitin before transfer to a substrate lysine, and the C4341A substitution abolishes this catalytic activity [19, 21]. Our results clearly demonstrated that the HUWE1(C4341A) mutant failed to promote K6-linked polyubiquitylation of Nrf2, confirming that catalytic activity is indispensable. Consistent with this, the small-molecule HUWE1 inhibitor BI8626, which targets the ubiquitin-binding site of the HECT domain [44], recapitulated the effects of HUWE1 knockdown on both Nrf2 ubiquitylation and HBV replication, validating the pharmacological approach.

A key finding of this study is that HUWE1 bound Nrf2 only in the presence of HBx, suggesting that HBx facilitates the association between HUWE1 and Nrf2. Under basal conditions, Nrf2 is degraded via Keap1-Cullin3-mediated K48-linked polyubiquitylation [8]. Our data support a model in which, during HBV infection, HUWE1 is recruited to Nrf2 regulatory context in an HBx-dependent manner, facilitating K6-linked modification associated with Nrf2 stabilization and suppressing HBV promoter activity. Further studies will be required to determine whether HBx acts as a direct molecular bridge and how this pathway is coordinated with canonical Keap1-dependent regulation.

HUWE1 has previously been reported to exhibit antiviral activity against other viruses. HUWE1 promotes ubiquitin-dependent degradation of MERS-CoV accessory protein ORF3 [45], negatively influences human immunodeficiency virus type 1 infectivity through interaction with viral Gag-Pol precursor protein [46], and acts as an antiviral E3 ligase against SARS-CoV-2 ORF9b [47]. Our study extends the antiviral repertoire of HUWE1 to HBV and suggests that HUWE1 may contribute to antiviral regulation in HBV infection by stabilizing a host transcription factor Nrf2 through non-canonical K6-linked polyubiquitylation.

Several limitations of this study should be acknowledged. First, although our data support the involvement of HUWE1 in HBx-dependent regulation of Nrf2, the precise lysine residue(s) on Nrf2 that receive K6-linked ubiquitin chains remain to be identified. Second, while our interaction data are consistent with a role for HBx in promoting HUWE1-Nrf2 complex formation, the molecular architecture of this complex and the potential contribution of additional host factors remain to be determined. Third, based on our previous finding that HBx interacts with Keap1, future studies should determine whether a complex containing HUWE1, HBx, Nrf2, and Keap1 is formed in HBV-infected cells and identify the HBx domains responsible for interaction with HUWE1 and Keap1. Finally, given that HUWE1 is involved in various cellular processes, including DNA damage response, autophagy, and apoptosis, through direct interactions with numerous proteins, such as p53, c-Myc, and Mcl-1 [12–14], we should examine the possibility that additional HUWE1-regulatory pathways also contribute to the antiviral phenotype observed here.

In conclusion, our study supports a model in which HUWE1 promotes HBx-dependent K6-linked polyubiquitylation and stabilization of Nrf2 and thereby contributes to the restriction of HBV replication. These findings expand current understanding of non-canonical ubiquitin signaling in HBV-host interactions and provide a basis for further mechanistic investigation.

## AUTHOR STATEMENTS

### Author Contributions

M.R.S., L.D., and I.S. conceived and designed the experiments. M.R.S. carried out most of the experiments. H.F.F., Y.P.K., C.M., and T.A. assisted with the construction and the data analysis. T.A., A.R., K.W., and M.M. contributed to the materials. M.R.S., L.D., and I.S. wrote the manuscript.

## Acknowledgments

We thank Y. Kozaki for her valuable secretarial assistance.

## REFERENCES

1. WHO. Global hepatitis report 2024: action for access in low- and middle-income countries. https://iris.who.int/server/api/core/bitstreams/4fc7b178-5017-4065-9970-fc5a7533cb5f/content (2024).

2. WHO. Hepatitis B. https://www.who.int/news-room/fact-sheets/detail/hepatitis-b (2025, accessed 4 March 2026).

3. Fanning GC, Zoulim F, Hou J, Bertoletti A. Therapeutic strategies for hepatitis B virus infection: towards a cure. Nat Rev Drug Discov 2019;18:827–844.

4. Lok ASF, McMahon BJ, Brown RS, Wong JB, Ahmed AT, et al. Antiviral therapy for chronic hepatitis B viral infection in adults: A systematic review and meta-analysis. Hepatology 2016;63:284–306.

5. Tsukuda S, Watashi K. Hepatitis B virus biology and life cycle. Antiviral Res 2020;182:104925.

6. Levrero M, Zucman-Rossi J. Mechanisms of HBV-induced hepatocellular carcinoma. J Hepatol 2016;64:S84–S101.

7. Tonelli C, Chio IIC, Tuveson DA. Transcriptional Regulation by Nrf2. Antioxid Redox Signal 2018;29:1727–1745.

8. Villeneuve NF, Lau A, Zhang DD. Regulation of the Nrf2–Keap1 Antioxidant Response by the Ubiquitin Proteasome System: An Insight into Cullin-Ring Ubiquitin Ligases. Antioxid Redox Signal 2010;13:1699–1712.

9. Hayes JD, Dayalan Naidu S, Dinkova-Kostova AT. Regulating Nrf2 activity: ubiquitin ligases and signaling molecules in redox homeostasis. Trends Biochem Sci 2025;50:179–205.

10. Kopacz A, Kloska D, Forman HJ, Jozkowicz A, Grochot-Przeczek A. Beyond repression of Nrf2: An update on Keap1. Free Radic Biol Med 2020;157:63–74.

11. Ariffianto A, Deng L, Abe T, Matsui C, Ito M, et al. Oxidative stress sensor Keap1 recognizes HBx protein to activate the Nrf2/ARE signaling pathway, thereby inhibiting hepatitis B virus replication. J Virol 2023;97:e01287–23.

12. Zhong Q, Gao W, Du F, Wang X. Mule/ARF-BP1, a BH3-Only E3 Ubiquitin Ligase, Catalyzes the Polyubiquitination of Mcl-1 and Regulates Apoptosis. Cell 2005;121:1085–1095.

13. Chen D, Kon N, Li M, Zhang W, Qin J, et al. ARF-BP1/Mule Is a Critical Mediator of the ARF Tumor Suppressor. Cell 2005;121:1071–1083.

14. Adhikary S, Marinoni F, Hock A, Hulleman E, Popov N, et al. The Ubiquitin Ligase HectH9 Regulates Transcriptional Activation by Myc and Is Essential for Tumor Cell Proliferation. Cell 2005;123:409–421.

15. Yoon SY, Lee Y, Kim JH, Chung A-S, Joo JH, et al. Over-expression of human UREB1 in colorectal cancer: HECT domain of human UREB1 inhibits the activity of tumor suppressor p53 protein. Biochem Biophys Res Commun 2004;326:7–17.

16. Liu Z, Oughtred R, Wing SS. Characterization of E3^Histone^ , a Novel Testis Ubiquitin Protein Ligase Which Ubiquitinates Histones. Mol Cell Biol 2005;25:2819–2831.

17. Kao S-H, Wu H-T, Wu K-J. Ubiquitination by HUWE1 in tumorigenesis and beyond. J Biomed Sci 2018;25:67.

18. Michel MA, Swatek KN, Hospenthal MK, Komander D. Ubiquitin Linkage-Specific Affimers Reveal Insights into K6-Linked Ubiquitin Signaling. Mol Cell 2017;68:233–246.e5.

19. Tracz M, Bialek W. Beyond K48 and K63: non-canonical protein ubiquitination. Cell Mol Biol Lett 2021;26:1.

20. Kulathu Y, Komander D. Atypical ubiquitylation — the unexplored world of polyubiquitin beyond Lys48 and Lys63 linkages. Nat Rev Mol Cell Biol 2012;13:508–523.

21. Hunkeler M, Jin CY, Ma MW, Monda JK, Overwijn D, et al. Solenoid architecture of HUWE1 contributes to ligase activity and substrate recognition. Mol Cell 2021;81:3468–3480.e7.

22. Suwardana GNR, Abe T, Deng L, Matsui C, Okitsu T, et al. A novel synthetic bile acid derivative inhibits hepatitis B virus infection at entry step by interfering with the oligomerization of sodium taurocholate co-transporting polypeptide. Antiviral Res 2025;243:106267.

23. Watashi K, Liang G, Iwamoto M, Marusawa H, Uchida N, et al. Interleukin-1 and Tumor Necrosis Factor-α Trigger Restriction of Hepatitis B Virus Infection via a Cytidine Deaminase Activation-induced Cytidine Deaminase (AID). J Biol Chem 2013;288:31715–31727.

24. Nishitsuji H, Ujino S, Shimizu Y, Harada K, Zhang J, et al. Novel reporter system to monitor early stages of the hepatitis B virus life cycle. Cancer Sci 2015;106:1616–1624.

25. Hayashi M, Deng L, Chen M, Gan X, Shinozaki K, et al. Interaction of the hepatitis B virus X protein with the lysine methyltransferase SET and MYND domain-containing 3 induces activator protein 1 activation. Microbiol Immunol 2016;60:17–25.

26. Sugiyama M, Tanaka Y, Kato T, Orito E, Ito K, et al. Influence of hepatitis B virus genotypes on the intra- and extracellular expression of viral DNA and antigens. Hepatology 2006;44:915.

27. Kouwaki T, Okamoto T, Ito A, Sugiyama Y, Yamashita K, et al. Hepatocyte Factor JMJD5 Regulates Hepatitis B Virus Replication through Interaction with HBx. J Virol 2016;90:3530–3542.

28. Fukutomi K, Hikita H, Murai K, Nakabori T, Shimoda A, et al. Capsid Allosteric Modulators Enhance the Innate Immune Response in Hepatitis B Virus–Infected Hepatocytes During Interferon Administration. Hepatol Commun 2022;6:281.

29. Deng L, Gan X, Ito M, Chen M, Aly HH, et al. Peroxiredoxin 1, a Novel HBx-Interacting Protein, Interacts with Exosome Component 5 and Negatively Regulates Hepatitis B Virus (HBV) Propagation through Degradation of HBV RNA. J Virol 2019;93:e02203–18.

30. Deng L, Liang Y, Ariffianto A, Matsui C, Abe T, et al. Hepatitis C Virus-Induced ROS/JNK Signaling Pathway Activates the E3 Ubiquitin Ligase Itch to Promote the Release of HCV Particles via Polyubiquitylation of VPS4A. J Virol;96:e01811–21.

31. Shirakura M, Murakami K, Ichimura T, Suzuki R, Shimoji T, et al. E6AP Ubiquitin Ligase Mediates Ubiquitylation and Degradation of Hepatitis C Virus Core Protein. J Virol 2007;81:1174–1185.

32. Benhenda S, Ducroux A, Rivière L, Sobhian B, Ward MD, et al. Methyltransferase PRMT1 Is a Binding Partner of HBx and a Negative Regulator of Hepatitis B Virus Transcription. J Virol 2013;87:4360–4371.

33. Ibrahim MK, Abdelhafez TH, Takeuchi JS, Wakae K, Sugiyama M, et al. MafF Is an Antiviral Host Factor That Suppresses Transcription from Hepatitis B Virus Core Promoter. J Virol 2021;95:10.1128/jvi.00767-21.

34. He Y, Zhou J, Wan Q. The E3 ligase HUWE1 mediates TGFBR2 ubiquitination and promotes gastric cancer cell proliferation, migration, and invasion. Invest New Drugs 2021;39:713–723.

35. Swatek KN, Komander D. Ubiquitin modifications. Cell Res 2016;26:399–422.

36. Yau R, Rape M. The increasing complexity of the ubiquitin code. Nat Cell Biol 2016;18:579–586.

37. Srivastava D, Chakrabarti O. Mahogunin-mediated α-tubulin ubiquitination via noncanonical K6 linkage regulates microtubule stability and mitotic spindle orientation. Cell Death Dis 2014;5:e1064–e1064.

38. Durcan TM, Tang MY, Pérusse JR, Dashti EA, Aguileta MA, et al. USP 8 regulates mitophagy by removing K 6-linked ubiquitin conjugates from parkin. EMBO J 2014;33:2473–2491.

39. Yuan Y, Miao Y, Qian L, Zhang Y, Liu C, et al. Targeting UBE4A Revives Viperin Protein in Epithelium to Enhance Host Antiviral Defense. Mol Cell 2020;77:734–747.e7.

40. Hong S-Y, Kao Y-R, Lee T-C, Wu C-W. Upregulation of E3 Ubiquitin Ligase CBLC Enhances EGFR Dysregulation and Signaling in Lung Adenocarcinoma. Cancer Res 2018;78:4984–4996.

41. Seo J, Lee E-W, Shin J, Seong D, Nam YW, et al. K6 linked polyubiquitylation of FADD by CHIP prevents death inducing signaling complex formation suppressing cell death. Oncogene 2018;37:4994–5006.

42. Zhang Z, Wang D, Wang P, Zhao Y, You F. OTUD1 Negatively Regulates Type I IFN Induction by Disrupting Noncanonical Ubiquitination of IRF3. J Immunol 2020;204:1904–1918.

43. Gong X, Du D, Deng Y, Zhou Y, Sun L, et al. The structure and regulation of the E3 ubiquitin ligase HUWE1 and its biological functions in cancer. Invest New Drugs 2020;38:515–524.

44. Peter S, Bultinck J, Myant K, Jaenicke LA, Walz S, et al. Tumor cell-specific inhibition of MYC function using small molecule inhibitors of the HUWE 1 ubiquitin ligase. EMBO Mol Med 2014;6:1525–1541.

45. Zhou Y, Zheng R, Liu S, Disoma C, Du A, et al. Host E3 ligase HUWE1 attenuates the proapoptotic activity of the MERS-CoV accessory protein ORF3 by promoting its ubiquitin-dependent degradation. J Biol Chem 2022;298:101584.

46. Yamamoto SP, Okawa K, Nakano T, Sano K, Ogawa K, et al. Huwe1, a novel cellular interactor of Gag-Pol through integrase binding, negatively influences HIV-1 infectivity. Microbes Infect 2011;13:339–349.

47. Yu M, Li J, Gao W, Li Z, Zhang W. Multiple E3 ligases act as antiviral factors against SARS-CoV-2 via inducing the ubiquitination and degradation of ORF9b. J Virol 2024;98:e01624–23.

